# rnaSPAdes: *a de novo* transcriptome assembler and its application to RNA-Seq data

**DOI:** 10.1101/420208

**Authors:** Elena Bushmanova, Dmitry Antipov, Alla Lapidus, Andrey D. Prjibelski

## Abstract

**Summary:** Possibility to generate large RNA-seq datasets has led to development of various reference-based and *de novo* transcriptome assemblers with their own strengths and limitations. While reference-based tools are widely used in various transcriptomic studies, their application is limited to the model organisms with finished and annotated genomes. *De novo* transcriptome reconstruction from short reads remains an open challenging problem, which is complicated by the varying expression levels across different genes, alternative splicing and paralogous genes. In this paper we describe a novel transcriptome assembler called rnaSPAdes, which is developed on top of SPAdes genome assembler and explores surprising computational parallels between assembly of transcriptomes and single-cell genomes. We also present quality assessment reports for rnaSPAdes assemblies, compare it with modern transcriptome assembly tools using several evaluation approaches on various RNA-Seq datasets, and briefly highlight strong and weak points of different assemblers.

**Availability and implementation:** rnaSPAdes is implemented in C++ and Python and is freely available at cab.spbu.ru/software/rnaspades/.

## 1 Introduction

While reference-based methods for RNA-Seq analysis [5,7,10,11,16,23] currently dominate transcriptome studies, they are subjected to the following constraints: (i) they are not applicable in the case when the genome is unknown, (ii) their performance deteriorates when the genome sequence or annotation are incomplete, and (iii) they may miss unusual transcripts (such as fusion genes or genes with short unannotated exons) even when the reference genome is available. To address these constraints, *de novo* transcriptome assemblers [6,15,19,20,25] have emerged as a viable complement to the reference-based tools. Although *de novo* assemblers typically generate fewer complete transcripts than the reference-based methods for the organisms with accurate reference sequences [12], they may provide additional insights on aberrant transcripts.

While the transcriptome assembly may seem to be a simpler problem than the genome assembly, RNA-Seq assemblers have to address the complications arising from highly uneven read coverage depth caused by variations in gene expression levels. However, this is the same challenge that we have addressed while developing SPAdes assembler [2, 14], which originally aimed at single-cell sequencing. Similarly to RNA-Seq, the Multiple Displacement Amplification (MDA) technique [8], used for genome amplification of single bacterial cells, results in a highly uneven read coverage. In the view of similarities between RNA-seq and single-cell genome assemblies, we decided to test SPAdes without any modifications on transcriptomic data. Surprisingly, even though SPAdes is a genome assembler and was not optimized for RNA-seq data, in some cases it generated decent assemblies of quality comparable to the state-of-the-art transcriptome assemblers.

To perform the benchmarking we have used rnaQUAST tool [4], which was designed for quality evaluation of *de novo* assemblies of model organisms, i.e. with the support of reference genome and its gene database. For the comparison, we have selected a few representative metrics such as (i) the total number of assembled transcripts, (ii) the number of transcripts that do not align to the reference genome, (iii) reference gene database coverage, (iv) the number of 50% / 95%-assembled genes/isoforms and (v) duplication ratio. The detailed description for these metrics can be found in the Supplementary material.

Table 1 demonstrates the comparison between Trans-ABySS [19], IDBA-tran [15], SOAPdenovo-Trans [25], Trinity [6] and SPAdes [2] on publicly available mouse RNA-Seq dataset. Trinity and IDBA-tran were launched with default parameters, Trans-ABySS and SOAPdenovo-Trans were run with *k*-mer sizes set to 32 and 31 respectively (see more information on selecting optimal *k*-mer lengths in the Results section), SPAdes was run in single-cell mode due to the uneven coverage depth of RNA-Seq data. Table 1 shows that SPAdes generates more 50% / 95%-assembled genes than any other assemblers and performs relatively well according to other parameters, such as database coverage. At the same time, SPAdes produces the highest number of misassembled transcripts, which can be explained by the fact that algorithms for genome assembly tend to assemble longer contigs and may incorrectly join sequences corresponding to different genes when working with RNA-Seq data. In addition, SPAdes generates exactly the same number 95%-assembled genes and isoforms, which emphasizes the absence of isoform detection step.

**Table 1:** Benchmarking of Trans-**ABySS**, **IDBA**-tran, **SOAP**denovo-Trans, **Trinity**, and **SPAdes** on *M. musculus* RNA-seq dataset (accession number SRX648736, 11 million Illumina 100 bp long paired-end reads). The annotated transcriptome of *M. musculus* consists of 38924 genes and 94545 isoforms. The best values for each metric are highlighted with bold.

| Assembler | ABySS | IDBA | SOAP | Trinity | SPAdes |
| --- | --- | --- | --- | --- | --- |
| Assembled transcripts | 63871 | 38304 | 61564 | 47717 | 48876 |
| Unaligned transcripts | 232 | 98 | 273 | 160 | 817 |
| Misassemblies | 156 | 272 | **35** | 247 | 456 |
| Database coverage, % | 17.7 | 16.9 | 17.1 | **18.4** | 17.9 |
| Duplication ratio | 1.09 | **1.004** | 1.013 | 1.155 | 1.015 |
| 50%-assembled genes | 6368 | 6562 | 6383 | 6695 | **6972** |
| 95%-assembled genes | 1763 | 1572 | 1804 | 2251 | **2391** |
| 50%-assembled isoforms | 6984 | 6795 | 6592 | **7461** | 7140 |
| 95%-assembled isoforms | 1815 | 1572 | 1818 | **2388** | 2391 |

Benchmarking on other datasets also showed that SPAdes successfully deals with non-uniform coverage depth and produces relatively high number of 50% / 95%-assembled genes in most cases. However, it also confirmed the problem of large amount of erroneous transcripts as well as relatively low number of fully reconstructed alternative isoforms in SPAdes assemblies. Based on the obtained statistics we have decided to adapt current SPAdes algorithms for RNA-Seq data with the goal to improve quality of the generated assemblies and develop a new transcriptomic assembler called rnaSPAdes. In this manuscript we describe major pipeline modifications as well as several algorithmic improvements introduced in rnaSPAdes that allow to avoid misassemblies and obtain sequences of alternatively spliced isoforms.

To perform excessive benchmarking of rnaSPAdes and other transcriptome assemblers mentioned above, among publicly available data we selected several RNA-Seq datasets sequenced from the organisms with various splicing complexity. For the generated assemblies we present quality assessment reports obtained with different *de novo* and reference-based evaluation approaches. In addition, based on these results we discuss superiorities and disadvantages of various assembly tools and provide insights on their performance.

## 2 Methods

Most of the modern *de novo* genome assembly algorithms for short reads rely on the concept of the de Bruijn graph [17]. While the initial study proposed to look for an Eulerian path traversing the de Bruijn graph in order to reconstruct genomic sequences, it appeared to be rather impractical due to the presence of complex genomic repeats and sequencing artifacts, such as errors and coverage gaps. Instead, genome assemblers implement various heuristic approaches, most of which are based on coverage depth, graph topology and the fact that the genome corresponds to one or more long paths traversing through the graph [14,26]. Indeed, the later observation is not correct for the case of transcriptome assembly, in which RNA sequences correspond to numerous shorter path in the graph. Thus, to enable high-quality assemblies from RNA-Seq data the majority of procedures in the SPAdes pipeline have to undergo major alternations.

SPAdes pipeline for genome assembly consists of the following major steps: (i) sequencing error correction using BayesHammer module [13], (ii) construction of the condensed de Bruijn graph, (iii) graph simplification, which implies removing chimeric and erroneous edges, and (iv) repeat resolution and scaffolding with exSPAnder algorithm [18, 24]. While BayesHammer works well on the data with highly uneven coverage depth and requires no change for RNA-Seq datasets, graph simplification and repeat resolution procedures strongly rely on the properties of genomic sequences and thus require significant modifications and novel functionality for *de novo* transcriptome assembly.

### 2.1. Simplification of the de Bruijn graph

During the graph simplification stage erroneous edges are removed from the de Bruijn graph based on various criteria in order to obtain clean graph containing only correct sequences (further referred to as an *assembly graph*). In the SPAdes pipeline the simplification process includes multiple various procedures that can be classified into three types: (i) trimming *tips* (dead-end or dead-start edges), *(ii)* collapsing *bulges* (alternative paths) and (iii) removing *erroneous connections* (chimeric and other false edges). Below we describe the key modifications introduced into rnaSPAdes simplification algorithms. We also provide comparison between initial and improved simplification procedure on several RNA-Seq datasets in the Supplementary material (Table 1).

#### 2.1.1 Trimming tips

In the de Bruijn graph constructed from DNA reads the major fraction of tips (edges starting or ending at a vertex without other adjacent edges) typically correspond to sequencing errors and thus have to be removed. Since, only a few tips are correct and either represent chromosome ends or formed by coverage gaps, the existing genome assemblers implement rather aggressive tip clipping procedures [2, 26] assuming that coverage gaps appear rather rarely. However, in the de Bruijn graph built from RNA-Seq data a significant amount of tips correspond to transcripts’ ends and thus have to be preserved. In order to keep correct tips and obtain full-length transcripts, rnaSPAdes uses lower coverage and length thresholds for tip trimming procedure than SPAdes (see details below).

In some cases, tips originated due to the sequencing errors in multiple reads from highly-expressed isoforms may have coverage above the threshold. While genome assemblers may also exploit relative coverage cutoff to remove such tips, in transcriptome assembly this approach may result in trimming correct tips corresponding to the ends of low-expressed isoforms. However, erroneous tips typically have a small difference from the correct sequence without errors (e.g. 1-2 mismatches). To address this issue, we align tips to the alternative (correct) edges of the graph (Fig. 1a) and trim them if the identity exceeds a certain threshold (similar procedure is implemented in truSPAdes, which was designed for True Synthetic Long Reads assembly [3]). In case when two tips correspond to the ends of an alternatively spliced isoforms, it is highly unlikely for them to have similar nucleotide sequences (Fig. 1b).

**Figure 1:**
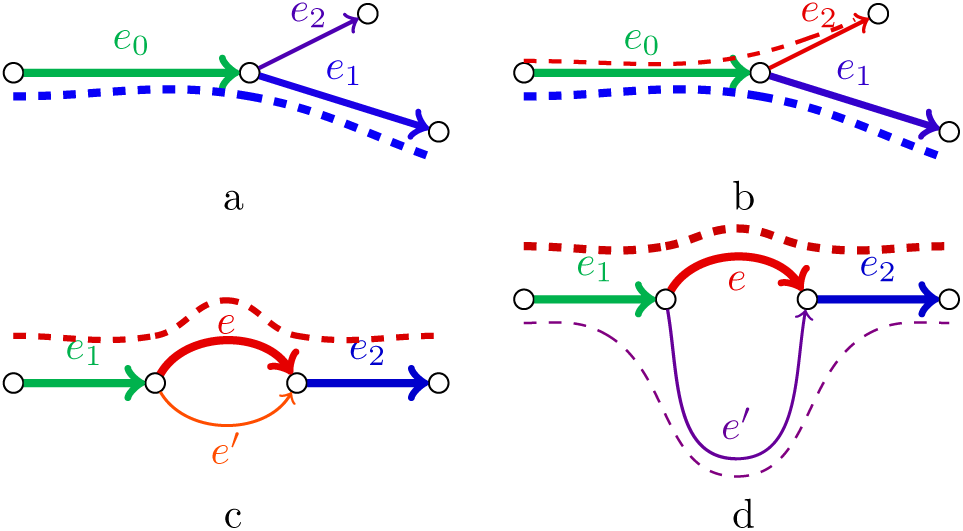
Examples of tips and bulges in the condensed de Bruijn graph. Edges with similar colors have similar sequences; line width represents the coverage depth. (a) Correct transcript (blue dashed line) traverses through edges *e*_0_ and *e*_1_. Edge *e*_2_ is originated from the reads with the same sequencing error and thus has coverage depth high enough not to be trimmed. However, since the sequence of edge *e*_2_ is very similar to the sequence of the alternative edge *e*_1_ (detected by alignment), *e*_2_ is eventually removed as erroneous. (b) In this case both paths (*e*_0_, *e*_1_) and (*e*_0_, *e*_2_) correspond to correct isoforms (blue and red dashed lines). Since the sequences of *e*_1_ and *e*_2_ are likely to be different, none of the correct tips is removed. (c) Correct sequence (red dashed line) traverses through edges *e*_1_, *e* and *e*_2_. Edge *e*^*′*^ is originated from reads containing sequencing errors, and thus has sequence similar to *e*, but significantly lower coverage. (d) Both paths (*e*_1_, *e, e*_2_) and (*e*_1_, *e*^*′*^, *e*_2_) correspond to different isoforms of the same gene (red and purple dashed lines); edges *e* and *e*^*′*^ typically have different length, coverage depth and sequence.

Another specifics of RNA-seq datasets is the large number of low-complexity regions that originate from poly-A tails resulting from polyadenylation at the ends of mRNAs. In order to avoid chimeric connections and non-informative sequences, we also remove low-complexity edges from the de Bruijn graph.

Below we summarize all conditions used in tip clipping procedure, parameters for which were optimized based on our analysis of various RNA-seq datasets. We define *l*_*T*_ as the length of the tip that is being analyzed and *c*_*T*_ as its mean *k*-mer coverage, and *c*_*A*_ as the *k*-mer coverage of the alternative edge (which is presumably correct) A tip is removed if any of the following conditions is true:

- *l* < 2 · *k* and *c*_*T*_ ≤ 1 (short tips with very low coverage);
- *l* < 4 · *k, c*_*T*_ < *c*_*A*_/2 and the Hamming distance between the tip and the alternative edge does not exceed 3 (the tip containing a sequencing error); the tip contains more than 80% of A nucleotides (T for reverse-complement edges).

#### 2.1.2 Collapsing bulges

A simple bulge (two edges sharing starting and terminal vertices) in the de Burijn graph may correspond to one of the following events: (i) a sequencing error, (ii) a heterozygous mutation or another allele difference or (iii) an alternative splicing event (typical for transcriptomic data). The first two cases are characterized by the bulge edges having similar lengths and sequences. However, edges formed by sequencing errors are typically short and have significantly different coverage depth, since it is unlikely for the same error to occur numerous times at the same position (Fig. 1c). Vice versa, in the case of allele difference bulge edges usually have similar coverage. Thus, genome assembly algorithms for bulge removal typically rely on the coverage depth [2, 26].

Since the most typical difference between two alternatively spliced isoforms of the same gene is the inclusion/exclusion of a an exon (usually short), edges of the bulge originated from these isoforms have different lengths (Fig. 1d). At the same time, since the expression levels may vary for such isoforms, the coverage depth may significantly differ. To avoid missing alternatively spliced isoforms in the assembly, rnaSPAdes does not use any coverage threshold for bulge removal and collapses only bulges consisting of edges with the similar lengths (less than 10% difference in length).

#### 2.1.3 Removing chimeric connections

While undetected tips and bulges formed by sequencing errors result in mismatches and indels in the assembled contigs, chimeric reads (typically corresponding to a concatenation of sequences from distant regions of the original molecules) may trigger more serious errors, such as incorrect junctions in the resulting contigs (often referred to as misassemblies). In conventional genome assembly chimeric edges usually have low coverage and thus can be easily identified [26]. Single-cell datasets, however, feature multiple low-covered genomic regions and elevated number of chimeric reads, which result in numerous erroneous connections having higher coverage depth than correct genomic edges. Similarly, since true edges representing low-expressed isoforms in the transcriptome assembly also have relatively low coverage depth, cleaning the graph using coverage threshold will result in multiple missing transcripts in the assembly.

To detect chimeric connections in single-cell assemblies SPAdes implements various algorithms, which mostly rely on the assumption that each chromosome corresponds to a long contiguous path traversing through the de Bruijn graph [14]. Since this assumption does not hold for transcriptomes consisting of thousands isoforms, we had to disable most procedures for the chimeric edge detection in SPAdes and implement a new erroneous edge removal algorithm that addresses the specifics of chimeric reads in RNA-seq data sets.

Our analysis revealed that most of the chimeric connections in RNA-seq data can be divided into two groups: single-strand chimeric loops and double-strand hairpins. In the first case, a chimeric junctions connects the end of a transcript sequence with itself (Fig. 2a). The erroneous hairpin connects correct edge with its reverse-complement copy (Fig. 2b) and potentially may result in chimeric palindromic sequence in the assembly. To avoid misassemblies, rnaSPAdes detects chimeric loops and hairpins by analyzing the graph topology rather than nucleotide sequences or coverage.

**Figure 2:**
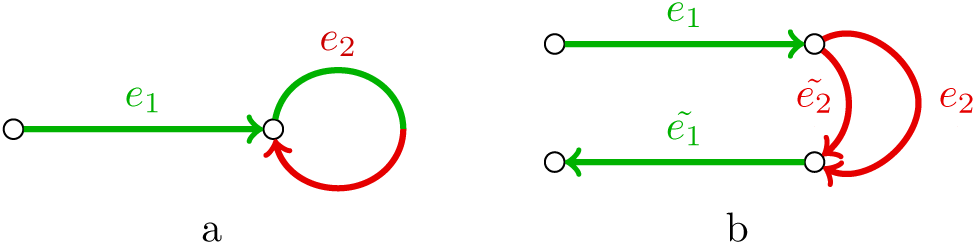
Examples of chimeric connections in the de Bruijn graph typical for transcriptome assembly. Red and green indicate erroneous and correct sequences respectively. (a) A chimeric loop (edge *e*_2_) connecting end of the correct transcriptomic edge *e*_1_ with itself. (b) An example of chimeric hairpin, where erroneous edge *e*_2_ connects a correct edge *e*_1_ with its reverse-complement copy *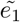*. Since *e*_2_ connects a vertex and its reverse-complement, *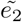* (the reverse-complement of *e*_2_) also connects these two vertices.

While it remains unclear whether these chimeric reads are formed during transcription, RNA-seq sample preparation or sequencing, similar chimeric connections have been observed in the context of single-cell MDA. E.g., when a DNA fragment is amplified by MDA, the DNA polymerase moves along DNA molecule and copies it, but sometimes (as described in [8]), the polymerase may jump to a close position (usually on the opposite DNA strand) and proceed to copy from the new position.

#### 2.1.4 Selecting optimal k-mer size

One of the key techniques that allows SPAdes to assemble contiguous genomic sequences from the data with non-uniform coverage depth is the iterative de Bruijn graph construction. During each next iteration, SPAdes builds the graph from the input reads and sequences obtained at the previous iteration, simplifies the graph and provides its edges as an input to the next iteration that uses larger *k*-mer size. Assembly graph obtained at the final iteration is used for repeat resolution and scaffolding procedures, which exploit read-pairs and long reads [1, 18]. In this approach, small *k*-mer sizes help to assemble low-covered regions where reads have short overlaps, and large *k* values are useful for resolving repeats and therefore obtaining less tangled graph. Although this method seems to be potentially useful for restoring low-expressed isoforms from RNA-Seq data, our analysis revealed that it appears to be the main reason of the high number of misassembled contigs in SPAdes assemblies (Table 1). Below we describe how these false junctions are formed.

When two transcripts (possibly from different genes) have a common sequence in the middle, they form a typical repeat structure in the de Bruijn graph (Fig. 3a), which may further be resolved, e.g. using paired reads. However, if a common sequence appears close to the ends of the transcripts (Fig. 3b), edges *e*_2_ and *e*_3_ appear to be rather short and may be trimmed as tips (since coverage depth often drops near the transcripts ends), or may not be present at all. In this case, the remaining edges *e*_1_, *e* and *e*_4_ will be condensed into a single edge corresponding to chimeric sequence.

**Figure 3:**
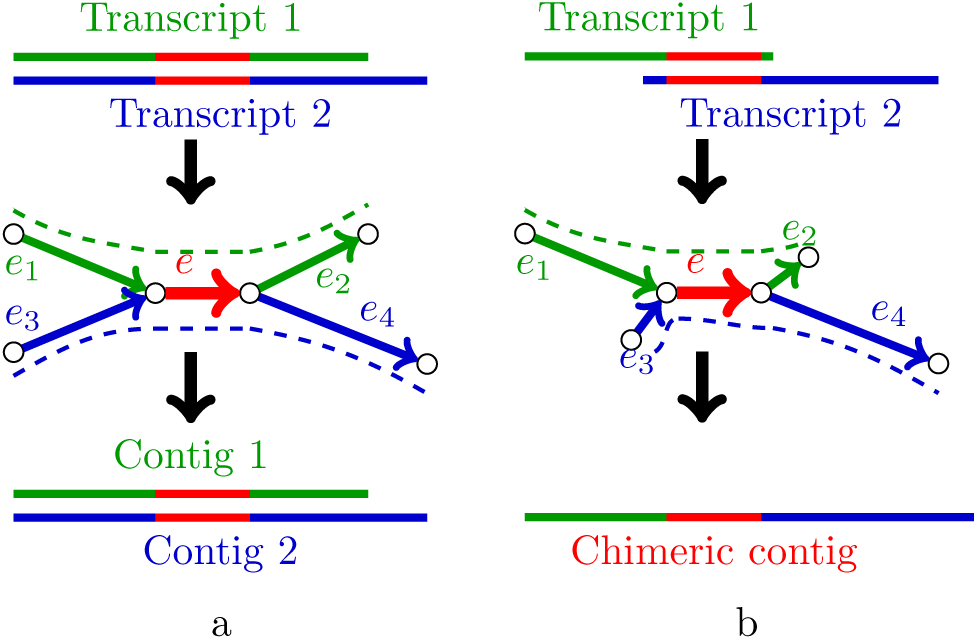
Examples of two transcripts having a common sequence (a) in the middle of the transcripts and (b) close to the start of one transcript and the end of another. While in the first case the isoforms can be resolved using read-pairs, the latter one may potentially result in a chimeric contig.

Indeed, since smaller *k*-mer size results in a higher chance of creating such kind of chimeric junction, we decided to disable the iterative graph construction and switch to a conventional approach of building a graph using a single large *k*-mer size (see Table 2 in the Supplementary material for comparison). To estimate the optimal *k* value, we ran rnaSPAdes on several RNA-Seq datasets with various read lengths sequenced from organisms with different gene complexity, and selected the best assemblies according to the number of assembled genes, the database coverage and the number of misassemblies. Although it may be not possible to choose a single best *k* value simultaneously for multiple datasets, nearly optimal *k*-mer size was estimated as the largest odd number that does not exceed *read_length/*2 – 1. All rnaSPAdes assemblies presented in this study are obtained using this default *k*-mer length, which is automatically calculated prior to the graph construction.

**Table 2:** Benchmarking of Trans-**ABySS** (*k* = 28), **IDBA**-tran, **SOAP**denovo-Trans (*k* = 25), **Trinity**, and **rnaSPAdes** (*k* = 43) on *C. elegans* non-strand-specific RNA-seq dataset (accession number SRR1560107, 9 million Illumina 90 bp long paired-end reads). The annotated transcriptome of *C. elegans* consists of 46748 genes and 57834 isoforms. The best values for each metric are highlighted with bold.

| Assembler | ABYSS | IDBA | SOAP | Trinity | rnaSPAdes |
| --- | --- | --- | --- | --- | --- |
| Transcripts | 64139 | 23157 | 28331 | 24450 | 33976 |
| Unaligned | 1 | **0** | 3 | 2 | 3 |
| Misassemblies | 46 | **27** | 50 | 153 | 57 |
| Database coverage | **39.2** | 33.2 | 34.1 | 36.2 | 36.6 |
| Duplication ratio | 1.659 | **1.007** | 1.023 | 1.177 | 1.08 |
| 50%-assembled genes | 10298 | 10090 | 10200 | 10423 | **10726** |
| 95%-assembled genes | 5434 | 3791 | 5417 | 5671 | **6326** |
| 50%-ass. isoforms | 11170 | 10244 | 10441 | **11185** | 11142 |
| 95%-ass. isoforms | 5500 | 3793 | 5517 | 6010 | **6523** |

In order to preserve correct connections that could be restored during the iterative graph construction, we carefully examined low-expressed transcripts that were not completely assembled using approach with a single *k*-mer size. The analysis revealed that the majority of such fragments can be joined by the small overlap, which is often confirmed by the read-pairs. To perform the gap closing procedure rnaSPAdes glues two tips if one of the following conditions is true:

- tips have an exact overlap of length at least *L*_*ov*_ and are connected by at least *N*_*ov*_ read pairs;
- tips are connected by at least *N*_*min*_ read pairs.

where the default parameters are *L*_*ov*_ = 8 bp, *N*_*ov*_ = 1 and *N*_*min*_ = 5. Although these parameters seem to be slightly ad-hoc, such gap closing procedure appears to be a viable alternative to the iterative graph construction and allows to restore more low-expressed transcripts without increasing the number of misassemblies. The analysis showed that while smaller thresholds often create false connections and increase the amount of erroneous transcripts, larger values for these parameters result in a smaller number of reconstructed sequences.

### 2.2. Isoform reconstruction

#### 2.2.1 Adapting repeat resolution algorithms

Genomic repeats present one of the key challenges in the *de novo* genome assembly problem. Although mRNA sequences typically do not contain complex repeats, transcriptome assembly has a somewhat similar problem of resolving alternatively spliced isoforms and isoforms from paralogous genes. Repeat resolution and scaffolding steps in SPAdes genome assembler are implemented in the exSPAnder module [18], which is based on simple path-extension framework. Similar to other modules of SPAdes, exSPAnder was designed to deal with highly uneven coverage and thus can be adapted for isoform detection procedure when assembling RNA-Seq data.

The key idea of the path-extension framework is to iteratively construct paths in the assembly graph by selecting the best-supported extension edge at every step until no extension can be chosen. The extension is selected based on the scoring function that may exploit various kinds of linkage information between edges of the assembly graph (different scoring functions are implemented for different types of sequencing data). A situation when a path cannot be extended further is usually caused by the presence of long genomic repeat or a large coverage gap. The extension procedure starts from the longest edge that is not yet included in any path and is repeated until all edges are covered.

More formally, a path extension step can be defined as follows. For a path *P* and its extension edges *e*_1_, *…, e*_*n*_ (typically, edges that start at the terminal vertex of *P*) the procedure selects *e*_*i*_ as a best-supported extension if

1. *Score*_*P*_ (*e*_*i*_) *> C · Score*_*P*_ (*e*_*j*_) for all *j ≠ i*

2. *Score*_*P*_ (*e*_*i*_) *>* Θ

where *C* and Θ are the algorithm parameters, and *Score*_*P*_ (*e*_*i*_) is a score of edge

*e*_*i*_ relative to path *P* (described in [18]).

In contrast to genome assembly, in which there is usually only one true extension edge, in transcriptome assembly multiple correct extensions are possible due to the presence of alternatively spliced isoforms. Thus, the modified procedure is capable of selecting several edges *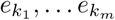* among all possible extensions *e*_1_, *…, e*_*n*_, which satisfy the following conditions:

1. *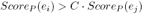* for all *i* = 1 *… m, M* = *argmax*_*j*=1..*n*_*Score*_*P*_ (*e*_*j*_)
2. *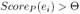* for all *i* = 1 *… m*

Namely, all correct extension edges must have a score close to the maximal one (*C* = 1.5 by default), and the second condition remains the same. Afterwards, the algorithm extends path *P* by creating new paths 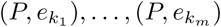, which are then extended independently. Since the scoring function implemented in exSPAnder does not strongly depend on the coverage depth, there is no danger that highly-expressed isoforms will be preferred over the low-expressed ones.

Finally, to avoid duplications in the genome assemblies, exSPAnder performs rather aggressive overlap removal procedure. However, since alternatively spliced isoforms may differ only by a short exon, in order to avoid missing similar transcripts the modified overlap detection procedure removes only exact duplicates and sub-paths.

#### 2.2.2 Exploiting coverage depth

Varying coverage depth may seem to be an additional challenge for *de novo* sequence assembly, but can be also used as an advantage in some cases. For instance, if two alternatively spliced isoforms of the same gene have different expression levels, they can be resolved using coverage depth even when the read-pairs do not help (e.g. shared exon is longer than the insert size). Although using coverage values becomes more complicated when a gene has multiple different expressing isoforms, our analysis of several RNA-Seq datasets revealed that such cases are rather rare and most of the genes have one or two expressing isoforms within a single sample.

To exploit the coverage depth we decided to add a simple, but reliable path-extension rule. Let the path *P* = (*e*_1_, *e*_2_, *e*_3_) have extension edges *e* and *e*^*′*^ (Fig. 4a), such that *cov*(*e*) *> cov*(*e* ^*′*^) and *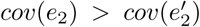*, where *cov*(*e*) denotes the *k*-mer coverage of edge *e*. To select a correct extension the algorithm detects a vertex closest to the end of path *P* that has two incoming alternative edges, one of which is included in *P* and another is not (*e*_2_ and *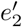* in this example). Since edge *e*_2_ ∊ *P* has higher coverage than the alternative edge *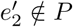*, we select extension edge *e* as the one with the higher coverage. However, if both isoforms have similar coverage, this simple approach may chose a false extension (since the coverage depth is rarely perfectly uniform even along a small region). Thus the difference in coverage should be significant enough to distinguish between the isoforms. More formally, the following conditions should be satisfied:

**Figure 4:**
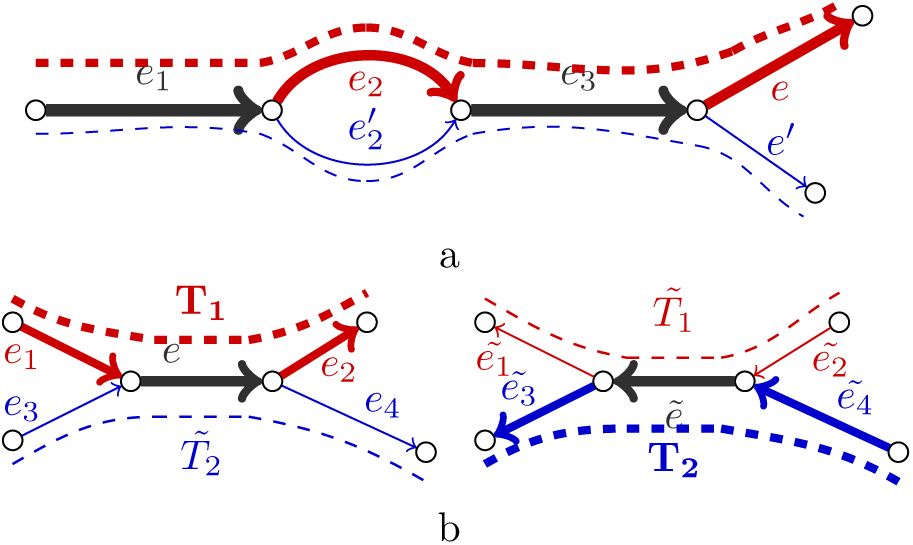
Using coverage depth for isoform reconstruction. Line width represents conventional and strand-specific coverage depths in figures (a) and (b) respectively. (a) Two isoforms of the same gene (red and blue dashed lines) have different expression levels and thus can be resolved using coverage depth. (b) Two transcripts *T*_1_ and *T*_2_ (red and blue bold dashed lines respectively) share a reverse-complement sequence and thus can be resolved using strand-specific reads.

1. *cov*(*e*) *>* Δ · *cov*(*e*^*′*^)
2. *cov*(*e*_2_) *>* Δ *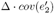*
3. Δ *> cov*(*e*_2_)*/cov*(*e*) *>* 1/Δ
4. *cov*(*e*) *> C*_*min*_

where the default values of the algorithm parameters are Δ = 2 and *C*_*min*_ = 2. The first two conditions ensure that the extension edges (*e* and *e*^*′*^) and alternative edges (*e*_2_ and *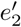*) have significant coverage difference, the third one requires the coverage depth to remain relatively persistent along the path and the latter one prevents the algorithm from resolving low-covered isoforms (which may result in a misassembly). In general case, this procedure also utilizes only the last pair of alternative edges and is applied only in case when the path has two possible extension edges and conventional read-pair extender fails to extend the path.

#### 2.2.3 Assembling strand-specific data

Another possible way to improve a transcriptome assembly is to take the benefit of strand-specific data when provided. To utilize stranded RNA-Seq we introduce *strand-specific coverage depths cov*^+^(*e*) and *cov*^-^(*e*), which denote *k*-mer coverage of edge *e* by forward and reverse reads respectively. As opposed to the conventional coverage *cov*(*e*), which is calculated by aligning all reads and their reverse-complement copies to the edges of the assembly graph (thus making *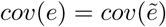*), strand-specific coverage is obtained by mapping reads according to their origin strand. For instance, if an RNA-Seq library is constructed in such way that reads have the same strand as the transcript which they were sequenced from, we expect *cov*^+^(*e*) to be much higher than *cov*^-^(*e*) if the sequence of *e* corresponds to the transcript, and vice versa if *e* is the reverse-complement of the original transcript. Indeed, the situation becomes opposite when reads are sequenced from cDNAs that are reverse-complement to the original transcripts. When working with paired-end libraries, we assume that the type of the library is defined by the first read’s strand. For example, if paired-end reads have common forward-reverse orientation, the second read in pair is reverse-complemented before mapping in order to match the strand of the first read.

To extend the paths we apply the same path-extension procedure described above for conventional coverage, but use strand-specific coverage values instead. Fig. 4b demonstrates a situation, when two transcript correspond to paths *T*_1_ = (*e*_1_, *e, e*_2_) and *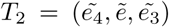*. If the repetitive edge *e* is longer than the insert size and the conventional coverage depth of these two transcripts is similar, the situation cannot be resolve neither by paired reads, nor by coverage. However, in case of stranded data, strand-specific coverage for actual transcripts’ paths will be much higher than for their reverse-complement copies, i.e. *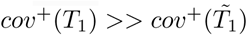* and *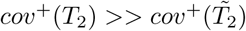* (in this example we assume that reads have the same stand as the transcripts they come from). Moreover, edges corresponding to the reverse-complement sequences only (*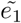* and *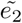* for 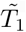, *e*_3_ and *e*_4_ for 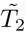) will have *cov*^+^(*e*) values close to zero. Therefore, the conditions given for coverage-based path extender (section 2.2.2) will be satisfied for strand-specific coverage values, the repetitive edge *e* will be resolved and both transcripts will be reconstructed.

In addition, for stranded RNA-Seq data we output the paths constructed by the exSPAnder algorithm according to the original transcript’s strand. E.g. in the example given in Figure 4b rnaSPAdes will output paths *T*_1_ and *T*_2_, since they have higher strand-specific coverage than their reverse complement copies (*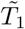* and *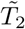* respectively).

#### 2.2.4 Filtering assembled transcripts

Before outputting the paths constructed by the exSPAnder module as contigs, we additionally apply various filtering procedures in order to remove non-mRNA contigs, such as intergenic sequences, which often contaminate RNA-Seq datasets. Our analysis showed that the majority of such unwanted sequences have low coverage, relatively small length and often correspond to isolated edges in the assembly graph (i.e. have no adjacent edges). However, applying filters based on these criteria may also remove correct low-expressed transcripts in some cases. Thus, we decided to implement three different presets of parameters for the filtration procedure (soft, normal and hard) and output three files with contigs. Depending on the project goal the researcher may choose more sensitive (soft filtration) or more specific results (hard filtration). Table 3 in the Supplementary material shows how the assembly quality depends on the filtration parameters. In other tables we use default transcripts with the normal level of filtering.

## 3 Results

### 3.1. Evaluating transcriptome assemblies for model organisms

To compare rnaSPAdes performance with the state-of-the-art transcriptome assemblers we selected several publicly available RNA-Seq datasets with different characteristics. In order to evaluate the resulting assemblies we used rnaQUAST [4], Transrate [22], BUSCO [21] and DETONATE package [9] (see details in the Supplementary material). In this manuscript, however, we present only statistics obtained with rnaQUAST for *C. elegans* assemblies (Table 2). Complete quality report for *C. elegans*, along with the results for *H. sapiens* and *M. musculus* non-stranded datasets and *Z .mays* strand-specific dataset can be also found in the Supplementary material (Tables 4-7 respectively).

To assemble selected datasets we launched IDBA-tran [15], Trinity [6] and rnaSPAdes with default parameters. Since *k*-mer size is a mandatory parameter and has no default value for Trans-ABySS [19] and SOAPdenovo-Trans [25], these tools were launched with the optimal *k*-mer sizes, which were selected individually for each dataset based on the number of assembled genes/isoforms, the amount of misassemblies and the database coverage. In order to make the results reproducible, we also provide software versions and command lines used in this study in the Supplementary material.

Table 2 demonstrates, that while all assemblies have comparable database coverage and approximately the same amount of 50%-assembled genes and isoforms, rnaSPAdes has the largest number of 95%-assembled genes and isoforms (11% and 8% more than the closest competitor — Trinity). While IDBA-trans assembly seems to be the most fragmented one, it has the smallest values of the duplication ratio and the number of misassemblies. According to these metrics both Trans-ABySS (highest duplication ratio) and Trinity (highest number of misassemblies) have less specific assemblies that contain redundant and erroneous sequences. However, despite the relatively high values for sensitivity metrics (50% / 95%-assembled genes/isoforms, database coverage), rnaSPAdes was able to maintain the appropriate levels of the duplication ratio (less than Trinity) and the number of misassemblies (comparable to SOAPdenovo-Trans and Trans-ABySS).

### 3.2 Discussions

Indeed, based on the provided benchmarks it is possible to highlight strong and weak points for every assembler included in the comparison. For example, Trinity and Trans-ABySS typically generate contigs with high duplication ratio and rather large number of misassemblies, which may negatively affect further *de novo* transcriptome analysis (such as annotation). In some cases, however, high values for these metrics may be caused by the presence of novel unannotated isoforms, which is most likely not the case for the high-quality reference genomes used in this study. At the same time both tools are rather sensitive and have decent database coverage and number of 50% / 95%-assembled genes/isoforms. Vice versa, IDBA-trans has the opposite assemblies’ characteristics, thus being the most specific and the least sensitive assembler in most cases. SOAPdenovo-Trans also generates rather correct assemblies with small number of erroneous transcripts, but has higher sensitivity metrics than IDBA-trans on average. Finally, rnaSPAdes always generates relatively high number of 50% / 95%-assembled genes, comparable values for database coverage and 50% / 95%-assembled isoforms, but at the same time manages to keep misassemblies and duplication ratio at moderate levels (lower than Trinity in all cases). These conclusions regarding specificity and sensitivity of different assemblies also correlate with the statistics reported by Transrate, BUSCO and DETONATE (Tables 4-7 in the Supplementary material).

Although every transcriptome assembler presented in this study has its own benefits and drawbacks, we believe that the trade-off between specificity and sensitivity can be significantly shifted by modifying the algorithms’ parameters. For example, various thresholds for transcript filtration in rnaSPAdes (Table 3 in the Supplementary material) result in assemblies with different properties (more specific assemblies for aggressive filtration and more sensitive for relaxed parameters). Also, incorporating iterative de Bruijn graph construction in rnaSPAdes (instead of building the graph with a single *k*-mer size) elevates the number of 50% / 95%-assembled genes/isoforms, but also raises the duplication ratio and the number of misassemblies (Table 2 in the Supplementary material). The same biases towards more sensitive or more specific results were observed during the selection of the optimal *k*-mer sizes for Trans-ABySS and SOAPdenovo-Trans. Thus, the parameters of the assembly algorithms can be varied in order to achieve the desired sensitivity or specificity characteristics and make the method to dominate by a certain group of metrics. In this paper, however, we mostly focus on rnaSPAdes modifications and parameters, rather than analyzing other assembly tools.

While the developed algorithm, rnaSPAdes, shows decent and stable results across multiple RNA-Seq datasets, the choice of the *de novo* transcriptome assembler remains a non-trivial problem, even with the aid of specially developed tools, such as Transrate, DETONATE, BUSCO and rnaQUAST. The selection of the assembler may be varied depending on the goals of the research project and the sample preparation protocols being used, as well as secondary parameters, such as usability and computational performance (see Table 8 in the Supplementary material for details).

#### Funding

The work was supported by Russian Science Foundation (grant 14-50-00069).

